# AGC Kinase Inhibitors Regulate STING Signaling Through SGK-Dependent and SGK-Independent Mechanisms

**DOI:** 10.1101/2022.07.21.500994

**Authors:** Johnny Castillo Cabrera, Hong Dang, Zhigang Zhang, Jose Torres-Castillo, Kelin Li, Pengda Liu, Jeff Aubé, Blossom Damania, Robert S. Hagan, Albert S. Baldwin

## Abstract

The STING signaling pathway is essential for the innate immune response to DNA viruses and bacteria and is important in tumor immunity. STING binding to cGAMP or to synthetic agonists leads to the activation of the kinase TBK1 which phosphorylates the transcription factor IRF3 which promotes expression of type 1 interferons such as IFNβ to block viral activity. Aberrant type 1 IFN expression is associated with human diseases including autoimmunity, HIV, and cancer. Here we identify N-[4-(1H-*pyrazolo*[3,4-b] pyrazin-6-yl)-phenyl]-sulfonamide (Sanofi-14h), a compound with preference for inhibition of the AGC family kinase SGK3, as an inhibitor of IFNβ gene expression in response to STING stimulation of macrophages. Sanofi-14h abrogated SGK activity and also impaired activation of the critical TBK1/IRF3 pathway downstream of STING activation, notably blocking the ligand-induced interaction of STING with TBK1. Deletion of SGK1 and SGK3 in macrophages suppressed activation of IFNβ transcription but did not block TBK1/IRF3 activation downstream of STING. Gene and protein expression analysis revealed that deletion of SGK1/3 in a macrophage cell line decreases basal expression of critical transcription factors required for the innate immune response, such as IRF7 and STAT1. Additional studies reveal that other AGC kinase inhibitors block TBK1 and IRF3 activation suggesting common action on a critical regulatory node in the STING pathway. Thus, studies with Sanofi-14h have revealed both SGK-dependent and SGK-independent effects in the STING pathway and suggest a mechanism to alter type 1 IFN transcription through small molecule therapy.

## Introduction

Type I interferons (IFNs) are cytokines that play critical roles in the innate immune response by inhibiting viral replication, promoting apoptosis of infected cells, recruiting and activating antigen presenting cells, and eliciting the adaptive immune response through activation of T and B cells^1^. However, dysregulated expression of interferons promotes autoimmune diseases^2^, inflammation^3^, and immunosuppressive stages of infection^4^. Underscoring the importance of interferons in the immune response, many pathogens express proteins to block interferon production leading to enhanced pathogenicity^5^. Understanding the regulation of IFN expression is critical for elucidating driving events and sequelae in infectious and chronic diseases, and for intervening therapeutically.

Expression of type I IFNs (α/β) is induced by stimulation of pathogen receptors including Toll-like receptors (TLR) and nucleic acid receptors such as cGAS^1, 6^. In the case of cGAS, binding to cytoplasmic DNA leads to the enzymatic generation of the cyclic dinucleotide 2’,3’ cGAMP^7^. Binding of bacteria- or cGAS-derived cyclic dinucleotides/cGAMP by STING leads to activation of the kinase TANK-binding kinase 1 (TBK1) and to the subsequent phosphorylation of key transcription factors interferon regulatory factor (IRF) 3 and 7, which directly promote transcription of type I IFN and other immune response genes^8–11^. Upregulated and secreted type I IFN signals through its receptor (IFNAR) to repress pathogen replication in infected and bystander cells and to activate transcription of a second wave of interferon-stimulated genes (ISGs). ISG expression downstream of IFN requires IFNAR-mediated activation of the JAK-STAT pathway with activating phosphorylation of the STAT1/STAT2/IRF9 (ISGF3) complex as well as STAT3 ^12, 13^ . Multiple layers of regulation by kinases, ubiquitin ligases, and other post-translational modifications tune both the baseline and activation state of the IFN pathway, representing an opportunity for development of chemicals that may repress or restore dysregulated IFN signaling.

Serum and glucocorticoid-regulated kinases (SGK1-3) are members of the AGC family of serine/threonine kinases with homology to the AKT subfamily^14^. SGK3 is widely expressed, whereas SGK2 expression is restricted largely to the pancreas, brain, kidneys and liver, and SGK1 is expressed in most tissues but with dynamic transcriptional and post-transcriptional regulation^15^. SGK3 uniquely contains a phosphoinositide-binding PX domain at the N-terminus which interacts with phosphatidylinositol-3-phosphate to control SGK3 localization at endosomes^16^. The activation of all three kinases is similar to that for AKT and is controlled by a priming phosphorylation by PI3K/PDK1 and activating phosphorylation by mTORC2^17^. The most well characterized SGK family substrates are members of the N-myc downstream regulated (NDRG) family in addition to several Na^+^/K^+^ ion channels ^18–22^. All three SGKs are linked to cancer either by promoting drug resistance or by controlling pathways important for cancer progression^15^. Bago et al demonstrated that PI3K/Akt inhibition in breast cancer cells leads to the upregulation of SGK3 to drive mTORC1 activity and tumor growth^23^, and Ranzuglia et al proposed SGK2 as regulator of autophagy that modulates platinum drug resistance in cancer cells^24^. SGK1 is the most characterized of the three in cancer with roles in driving tumorigenesis, proliferation, regulation of apoptosis, invasion, metastasis and drug resistance^25^.

SGK inhibition has been proposed as a therapeutic approach in cancer and SGK inhibitors have been developed and tested with this objective^26^. GSK650394, SI113 and EMD638683 are the most widely used inhibitors of SGK in cancer studies and have been shown to block cell proliferation and to decrease tumor growth alone or in combination with other inhibitors in animal models ^27–29^. A family of N-[4-(1H-*pyrazolo*[3,4-b] pyrazin-6-yl)-phenyl]-sulfonamides have been described to have more potent and selective inhibitory effects on SGK than previous inhibitors^30^. Furthermore, one derivative of these compounds, named Sanofi-14h, has been used in breast cancer models *in vivo* and *in vitro* and in combination with AKT inhibitors ^23^. Another derivative (Sanofi-17a) has been studied in a mouse explant model of osteoarthritis^31^. Notably, Sanofi-14h shows higher activity towards SGK3 as compared with SGK1, but also has significant activity against kinases such as S6K1 and MLK3^23^. Sanofi-17a preferentially inhibits SGK1 *in vitro* and has decreased activity against other AGC kinases including SGK3 and S6K^31^.

Studies on the functions of SGK kinases in innate immunity are limited. Reports suggest that SGK1 negatively regulates the TLR4/NF-κB response in mice ^32^, promotes macrophage polarization to an M1 phenotype^33^, and controls macrophage migration in vascular inflammation^34^. SGK3 was shown to regulate Ca^2+^-mediated dendritic migration in response to lipopolysaccharides (LPS) in mice^35^. SGK1 has been also reported to phosphorylate IKKα at Thr23 and activate NF-κB in Neuro2A cells and to phosphorylate IKKβ(S177A/S181A), but not IKKα, in breast cancer cells to increase NF-κB activity^36, 37^.

While testing compounds for effects on IFNβ expression we found that Sanofi-14h impairs IFNβ expression in macrophages downstream of STING. We show here that Sanofi-14h blocks the phosphorylation of TBK1 and IRF3, along with the dimerization and nuclear translocation of IRF3 following treatment with a STING agonist. Deletion of SGK1 and SGK3 in macrophages phenocopies the transcriptional defect of Sanofi-14h-treated cells. However, the SGK1/3-depleted cells retain TBK1 and IRF3 phosphorylation downstream of STING, and Sanofi-14h is able to block TBK1/IRF phosphorylation in SGK1/3-null cells, indicating both SGK-dependent and -independent effects in the response of macrophages to STING activation. Transcription and protein analysis revealed the effect of loss of SGK1 and SGK3 on IFNβ gene expression is through loss of basal expression of several key IRF and STAT family members. Interestingly, Sanofi-14h blocks the recruitment of TBK1 and IRF3 into the STING complex, indicating its ability to block a critical initial step in the STING-TBK1-IRF3 activation pathway. Analysis of other AGC kinase inhibitors across different drug classes revealed that many suppress IRF3 activation albeit with different efficacy. These studies indicate unexpected effects of AGC kinase inhibitors on the STING-driven innate immune response with a primary effect at the level of TBK1 and IRF3 activation. Additionally, these studies reveal regulatory effects of SGK1/3 in control of expression of critical transcription factors involved in the innate immune response.

## Results

### Sanofi-14h blocks IFNβ gene induction induced by STING ligands and HSV-1

While investigating compounds that may affect the induction of IFNβ gene transcription in the STING pathway, we found that N-[4-(1H-*pyrazolo*[3,4-b] pyrazin-6-yl)-phenyl]-sulfonamide (Sanofi-14h) suppressed IFNβ mRNA induction in murine RAW264.7 macrophages treated with DMXAA^38^ . The inhibitory response was dose-dependent with a 50% inhibition measured between 500nM and 1μM [**Figure 1A**]. To confirm the biological significance of this response, we analyzed the effect of Sanofi-14h on replication of the dsDNA herpes simplex virus (HSV-1). Consistent with **Figure 1A**, we found that Sanofi-14h increased replication of HSV-1 in RAW264.7 macrophages as measured through plaque-forming assays [**Figure 1B**]. Sanofi-14h also blocked induction of IFNβ mRNA in HSV-1 infected cells [**Figure 1B**].

**Figure 1:**
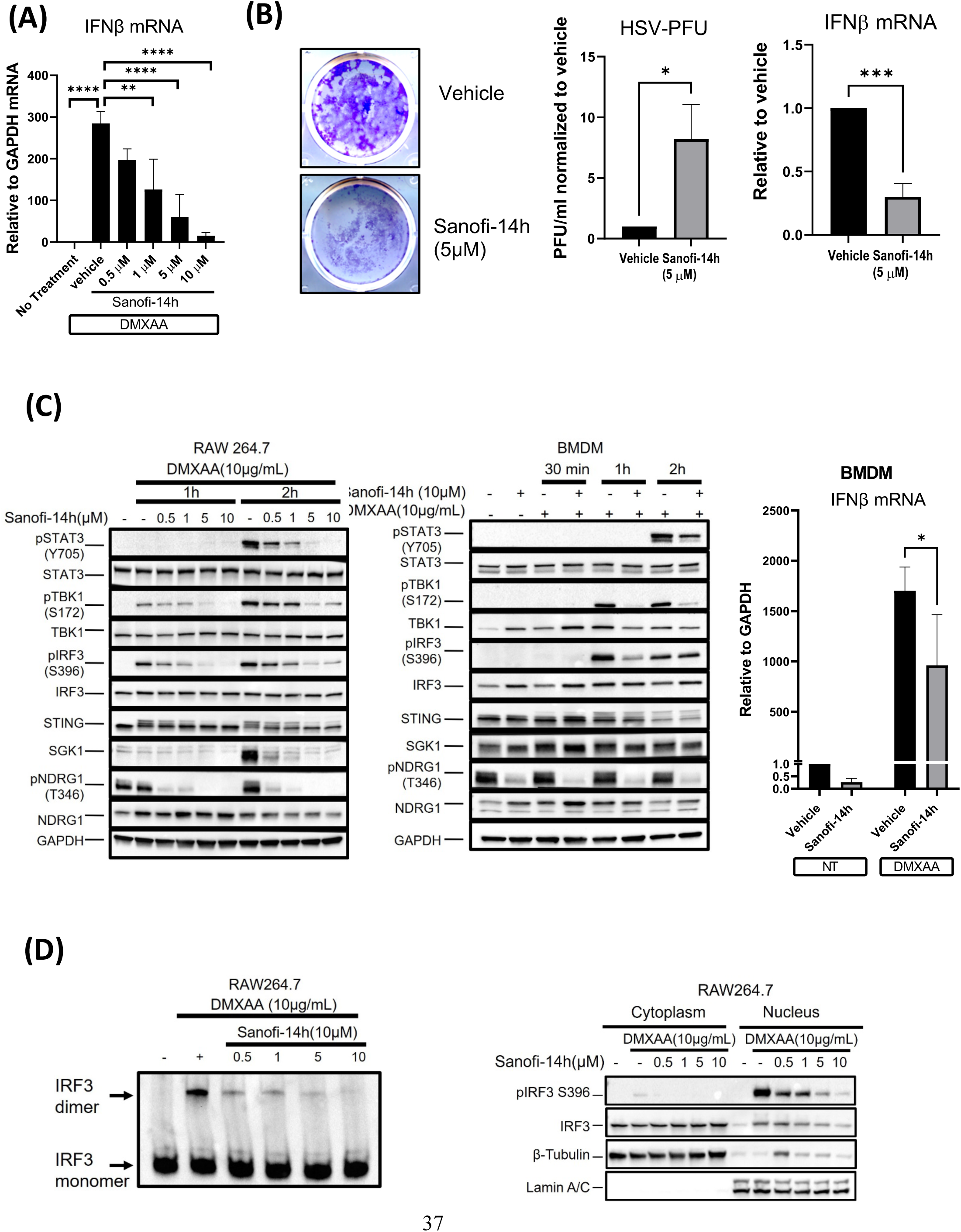
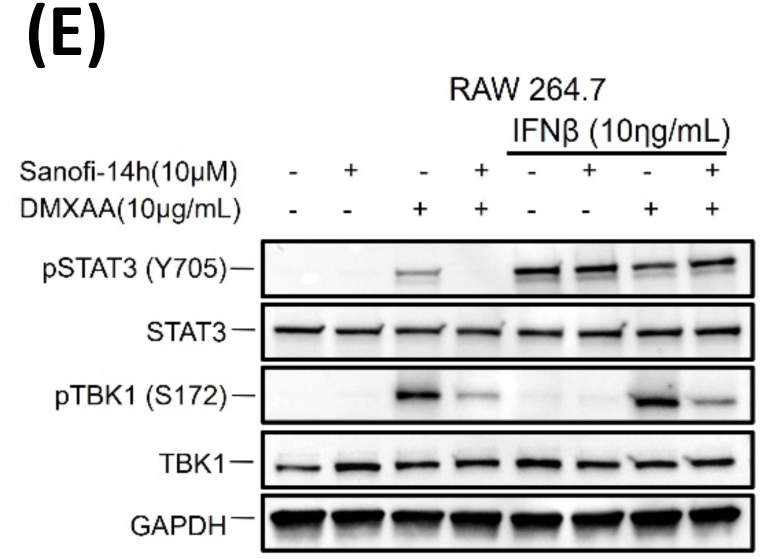
SGK inhibitor Sanofi-14h impairs IFN expression and STING/TBK1/IRF3 signaling in macrophages. **A)** RAW264.7 cells were pre-treated with Sanofi-14h at the indicated concentrations for 2 hours and then treated with DMXAA (10μg/mL) for an additional 2 hours. RT-qPCR IFNβ mRNA expression was normalized to the housekeeping gene GAPDH. **B)** Cells were seeded in 12 well plates and pre-treated the following day with Sanofi-14h (5 μM) for 24 h and then infected with HSV-1 KOS (MOI 0.1). The panel shows the plaque assay, and the graphs represents the PFU quantification (left) and FNβ mRNA after HSV-1 KOS infection(right). **C)** RAW264.7 macrophages (left panel) were plated in 60 mm plates and treated the next day with Sanofi-14h at the indicated doses for 2 h, followed by treatment with DMXAA (10μg/mL) for 1 and 2 h and immunoblotted. BMDM (right panel) were plated in the same way and treated with Sanofi-14h (10 μM) for 2 h and then with DMXAA (10μg/mL) for 30 min, 1 and 2 h. The graph shows the IFNβ mRNA expression normalized to GAPDH in BMDM. pNDRG1(T346) was a used a control to measure Sanofi-14h activity in both cell lines. **D)** Dimerization (left panel) and nuclear localization (right panel) of IRF3 after pre-treatment with Sanofi-14h at different doses for 2 hours and DMXAA for 1 hour. Lamin A/C and β-tubulin serve as control for the nuclear and cytoplasmic fractions, respectively. **E)** STAT3 phosphorylation under Sanofi-14h treatment, then with DMXAA and exogenous IFNβ treatment. After pretreatment with and without Sanofi-14h (10 μM) for 2 hours, cells were treated with DMXAA for either 2 h (control and control+Sanofi-14h) or 1 h and 45 min complemented with exogenous IFNβ (10 ng/mL) for 15 min. At the end all DMXAA samples were treated for a total of 2 h. BMDM: Bone Marrow Derived Macrophages, PFU: Plaque-Forming Unite, HSV: Herpes Simplex Virus. Data are representative of three different experiments (* p<0.05).

### Sanofi-14h blocks the STING-induced TBK1-IRF3 signaling cascade

STING activation in response to cytoplasmic DNA/viral infection leads to the phosphorylation of the kinase TBK1 which phosphorylates the transcription factor IRF3, leading to its subsequent dimerization and nuclear translocation to promote transcription of the IFNβ gene ^13, 39^. To determine whether Sanofi-14h affects this signaling pathway, we treated RAW264.7 cells with different doses of the inhibitor prior to stimulation with DMXAA. Sanofi-14h pre-treatment led to a dose-dependent inhibition of activating phosphorylation of TBK1, IRF3, and STING [**Figure 1C**]. STING phosphorylation is characterized by the reduction in its electrophoretic mobility. Consistent with the ability of Sanofi-14h to block induction of IFNβ, treatment with this inhibitor led to a marked decrease in STAT3 phosphorylation [**Figure 1C**], which occurs downstream of IFN receptor signaling. We observed that submicromolar doses of Sanofi-14h efficiently blocked phosphorylation of the canonical SGK target NDRG1, while higher doses of Sanofi-14h (>5 μM) potently impaired TBK1 and IRF3 phosphorylation [**Figure 1C**]. Both SGK1 protein [**Figure 1C**] and mRNA levels [**Suppl. Figure 1A**] are induced by DMXAA treatment and this increase is blocked in a dose-dependent manner by Sanofi-14h. These results indicate that SGK1 is a target gene downstream of STING activation.

Additionally, we assayed STING-induced signaling and IFNβ expression in primary mouse bone marrow-derived macrophages (BMDMs) treated with Sanofi-14h. Sanofi-14h efficiently blocks phosphorylation of NDRG1 in BMDMs [**Figure 1C**]. In addition, Sanofi-14h blocked induction of IFNβ mRNA and blocked activating phosphorylation of STAT3, TBK1, and IRF3 in primary macrophage cells [**Figure 1C**].

After phosphorylation, IRF3 transcription factor dimerizes and translocates to the nucleus to engage target genes for activation^39^. To further address downstream effects of Sanofi-14h on STING-induced signaling, we measured IRF3 dimerization and nuclear translocation in the presence of the drug following DMXAA stimulation of RAW264.7 cells. We observed that Sanofi-14h inhibited both dimerization and nuclear accumulation of IRF3 in a dose-dependent manner [**Figure 1D**]. Because STAT3 Y705 phosphorylation was blocked by Sanofi-14h, we asked whether this effect was dependent on type 1 IFN expression or via an intrinsic interaction between STAT3 and either SGK1/3 or TBK1. In this regard, TBK1 has been reported to phosphorylate STAT3 at S754 and to modulate its function in the STING pathway in macrophages^40^. We co-treated RAW264.7 cells with exogenous IFNβ, DMXAA and Sanofi-14h. We observed that treatment with exogenous IFNβ rescues the STAT3 phosphorylation inhibition in the presence of Sanofi-14h but does not rescue TBK1 phosphorylation inhibition induced by Sanofi-14h [**Figure 1E**]. This result indicates that the loss of p-STAT3 induced by Sanofi-14h is dependent on loss of IFNβ production.

### The effects of Sanofi-14h on TBK1/IRF3/STING activation are largely independent of SGK1 and SGK3

In RAW264.7 cells, SGK3 is expressed at relatively high levels and SGK1 is expressed at lower levels but is inducible upon STING stimulation [see **Figure 2D**, and **suppl.** **Fig 1A**]. SGK2 is not detected in these cells and is not upregulated with SGK1 or SGK3 knockdown [**Suppl. Figure 1B**]. To determine whether SGK1 and/or 3 are involved in control of IFNβ production and in STING-induced signaling, we generated single and double knockouts (KOs) of SGK1 and SGK3 in RAW264.7 cells using CRISPR-Cas9 [**Figure 2D**]. We then asked whether SGK1/3 KO cells phenocopy cells treated with Sanofi-14h prior to STING agonist treatment. First, we showed that loss of either SGK1 or SGK3 reduced IFNβ mRNA induction following DMXAA treatment [**Figure 2A**]. Next, we analyzed the effect of HSV1 infection on wild-type and SGK1/3 KO cells. RAW264.7 macrophages with single or double deletion of SGK1 or SGK3 showed increased cytopathic effect after HSV-1 infection [**Figure 2B**]. SGK1/3 KO macrophages showed an increase in viral proliferation and decreased IFNβ transcription. The effect of SGK3 deletion on IFNβ mRNA was more pronounced than SGK1 deletion, likely reflecting the higher baseline expression of SGK3 protein in RAW264.7 cells [**Figure 2B**]. We generated primary bone marrow-derived macrophages (BMDMs) from SGK3 null mice ^41^ and infected them with HSV *in vitro*. SGK3 null BMDMs showed increased viral replication as compared with wildtype cells [**Figure 2C**]. Thus, deletion of SGK1/3 in macrophages impairs *in vitro* antiviral defense similar to treatment with Sanofi-14h.

**Figure 2:**
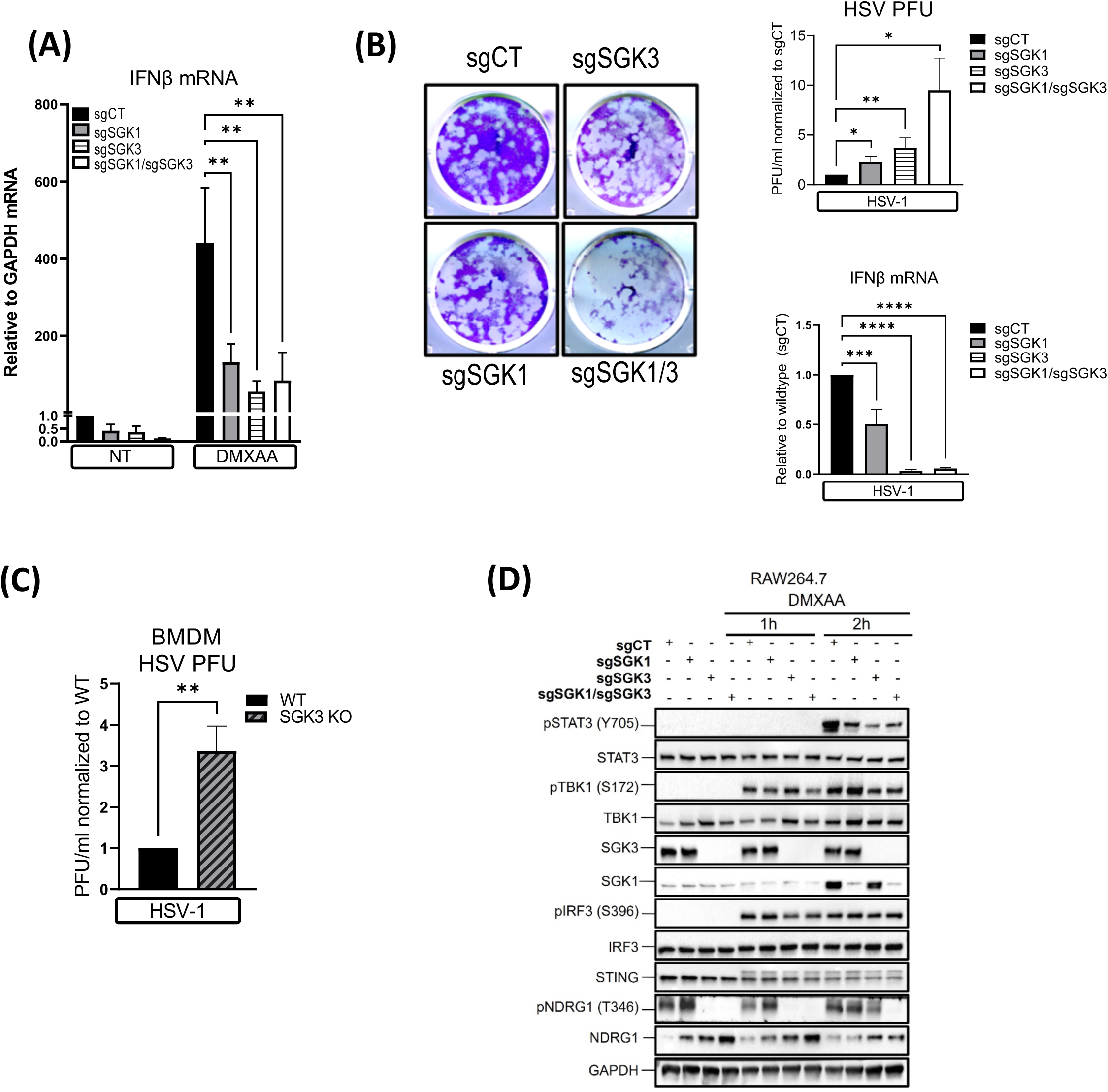
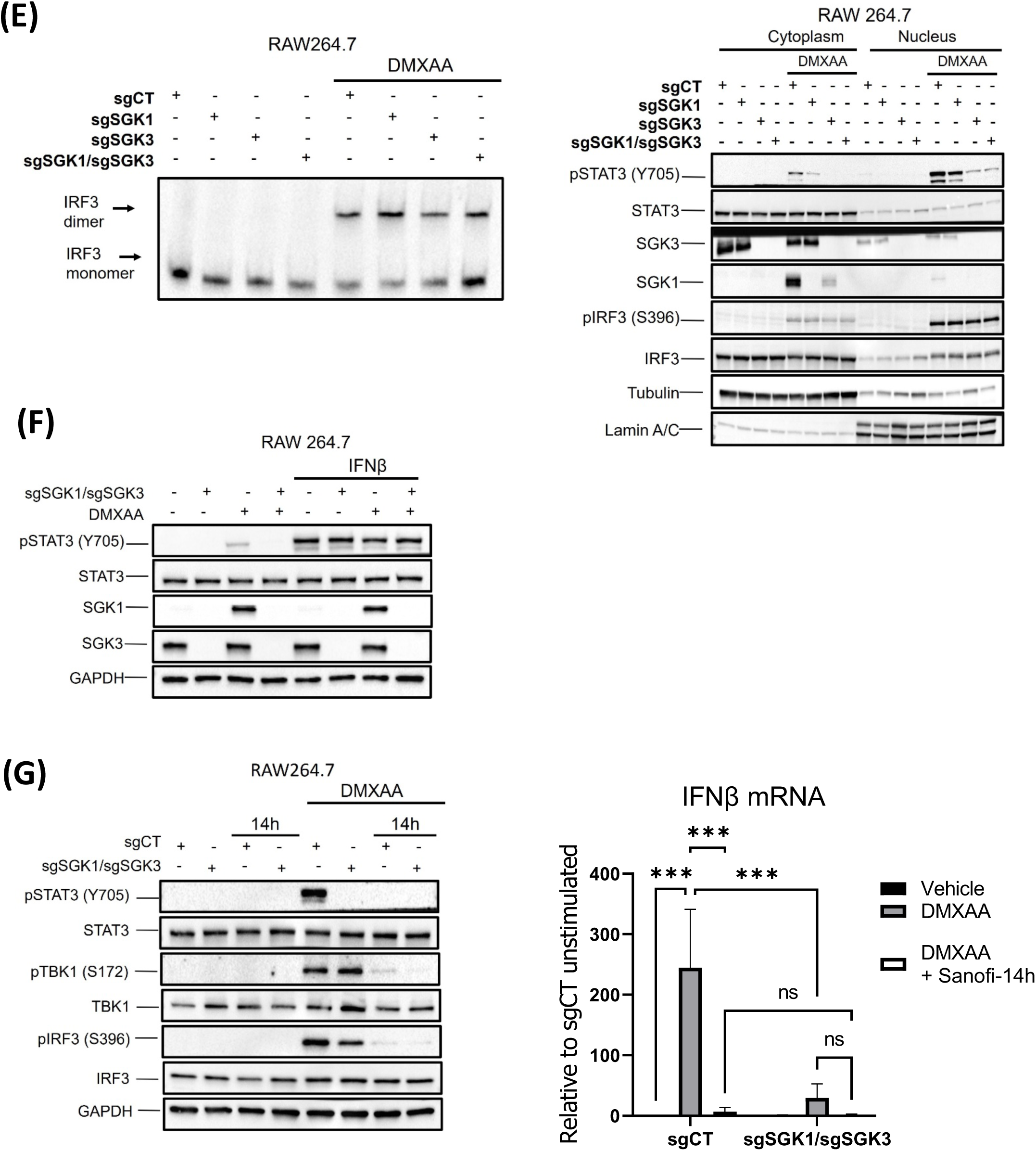
SGK1/3 KO does not affect TBK1, IRF3 and STING phosphorylation but hinders IFNβ production and consequential STAT3 phosphorylation. **A)** IFNβ mRNA expression in single and double(dKO) CRISPR generated RAW264.7 SGK1/3 KO cells with (top segment) and without (bottom segment) DMXAA treatment for 2 h. **B)** RAW264.7 SGK KO cells were plated in 12 well plates and infected with HSV-1 KOS. The upper panel shows the plaque assay for single and dKO cells, whereas the upper right graph represents the PFU and lower right graph shows IFNβ mRNA production in response to HSV infection these cells. **C)** PFU after HSV infection in BMDM SGK3 KO cells. **D)** Cells were treated with DMXAA for 1 and 2 h and then immunoblotted for different markers of activation. SGK1 and SGK3 protein levels serve as evidence of the knock out and pNDRG1 function as a control for lost activity. **E)** Dimerization (left panel) and nuclear localization (right panel) of IRF3 after DMXAA treatment for 1 h in SGK single and dKO cells. Lamin A/C and β-tubulin serve as control for the nuclear and cytoplasmic fractions, respectively. **F**) Immunoblot representing the treatment of SGK1/3 dKO cells with DMXAA (2 h total) with and without exogenous IFNβ (15 min). **G**) sgCT and sgSGK1/sgSGK3 cells were treated with Sanofi-14h for 2 h and the treated with DMXAA for an additional two hours before being immunoblotted for activation markers (left panel) and prepare for RT-qPCR (right graph) for IFNβ mRNA production. Data are representative of three different experiments (* p<0.05).

We next examined the signaling consequences of SGK deletion in response to DMXAA. We found that SGK3 KO cells demonstrate loss of basal NDRG1 phosphorylation, indicating that in RAW264.7 macrophages SGK3 is the primary NDRG1 kinase [**Figure 2D**]. SGK1 KO, SGK3 KO, and dual SGK1/3 KO cells exhibit decreased STAT3 Y705 phosphorylation in response to DMXAA treatment, in agreement with reduced IFNβ expression. Surprisingly, SGK1 KO, SGK3 KO, and double KO (dKO) cells have largely intact DMXAA-induced phosphorylation of TBK1 and STING [**Figure 2D**]. Consistent with this finding, SGK1/3 KO cells have largely unimpaired IRF3 phosphorylation after STING stimulation [**Figure 2D**]. DMXAA-stimulated dimerization of IRF3 is likewise unaffected by single or dual deletion of SGK1 or SGK3 [**Figure 2E**]. Fractionation of the nuclear and cytoplasmic compartments showed reduced nuclear accumulation of phosphorylated STAT3 but not IRF3 in SGK3 KO and SGK1/3 dKO cells [**Figure 2E**]. To test if SGK1/3 KO affects STAT3 through IFN expression or another mechanism, we treated WT and SGK1/3 dKO cells with exogenous IFNβ and found that IFNβ rescues the loss of STAT3 phosphorylation in the dKO cells, indicating that SGK1/3 activate STAT3 via control of IFNβ expression [**Figure 2E**].

As indicated in **Figure 2D**, loss of SGK1/3 did not block the phosphorylation of TBK1 or IRF3 downstream of DMXAA stimulation, yet Sanofi-14h treatment blocked activation of TBK1 as well as IRF3 in dKO cells [**Figure 2G**]. While knockout of SGK1/3 reduced IFNβ mRNA induction downstream of STING treatment, Sanofi-14h further reduced that response in these cells [**Figure 2G**]. Collectively, these data indicate that Sanofi-14h inhibition of TBK1/IRF3 phosphorylation during STING activation is largely independent of SGK1/3 inhibition and that SGK1/3 function through a distinct mechanism to promote STING-induced IFNβ mRNA production.

### SGK1 and SGK3 control baseline and inducible expression of innate immune regulators

We next performed RNA sequencing (RNAseq) on WT and SGK1/3 dKO RAW264.7 cells with and without DMXAA stimulation. Unsupervised clustering of unstimulated samples showed clear delineation of WT and SGK-null samples with a large number of genes dysregulated in SGK1/3 KO cells. Gene ontology analysis revealed basal downregulation of genes associated with pattern recognition receptor (PRR) signaling, innate immune response, interferon production, and responses to viral infection in cells lacking SGK1 and SGK3 [**Figure 3A**]. These SGK-dependent genes include IFN regulating transcription factors Stat1, Irf7, and Irf9. Volcano plots show SGK-dependent baseline expression of other innate immune genes and Interferon-Stimulated Genes (ISGs) including Ifi44, Il1rn, Usp14, Siglec1, and Oas3.Gene set enrichment analysis (GSEA) comparing dysregulated genes in WT and SGK1/3 dKO cells at baseline indicated loss of gene sets related to host defense with no other strongly significant gene sets identified.

**Figure 3:**
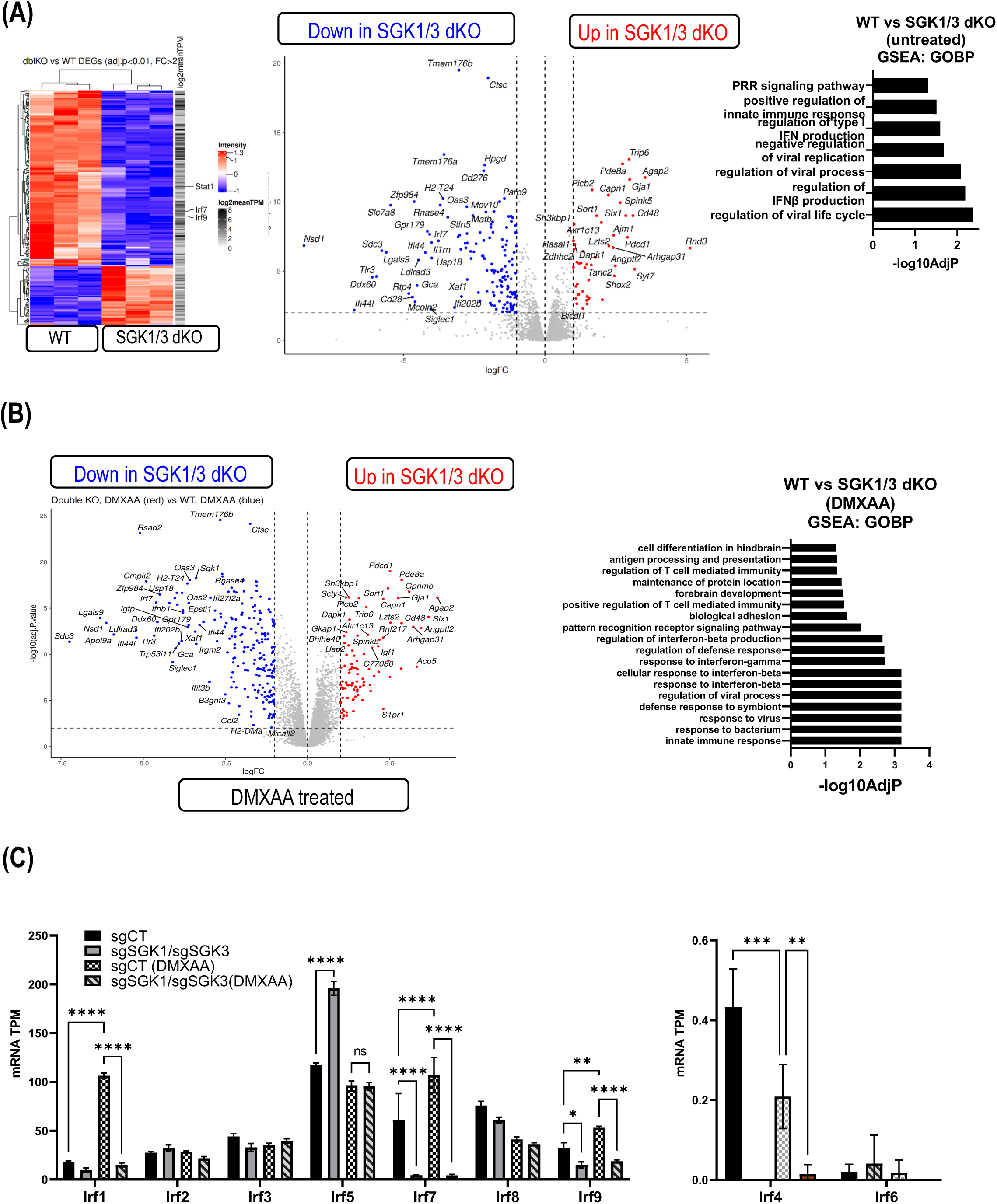
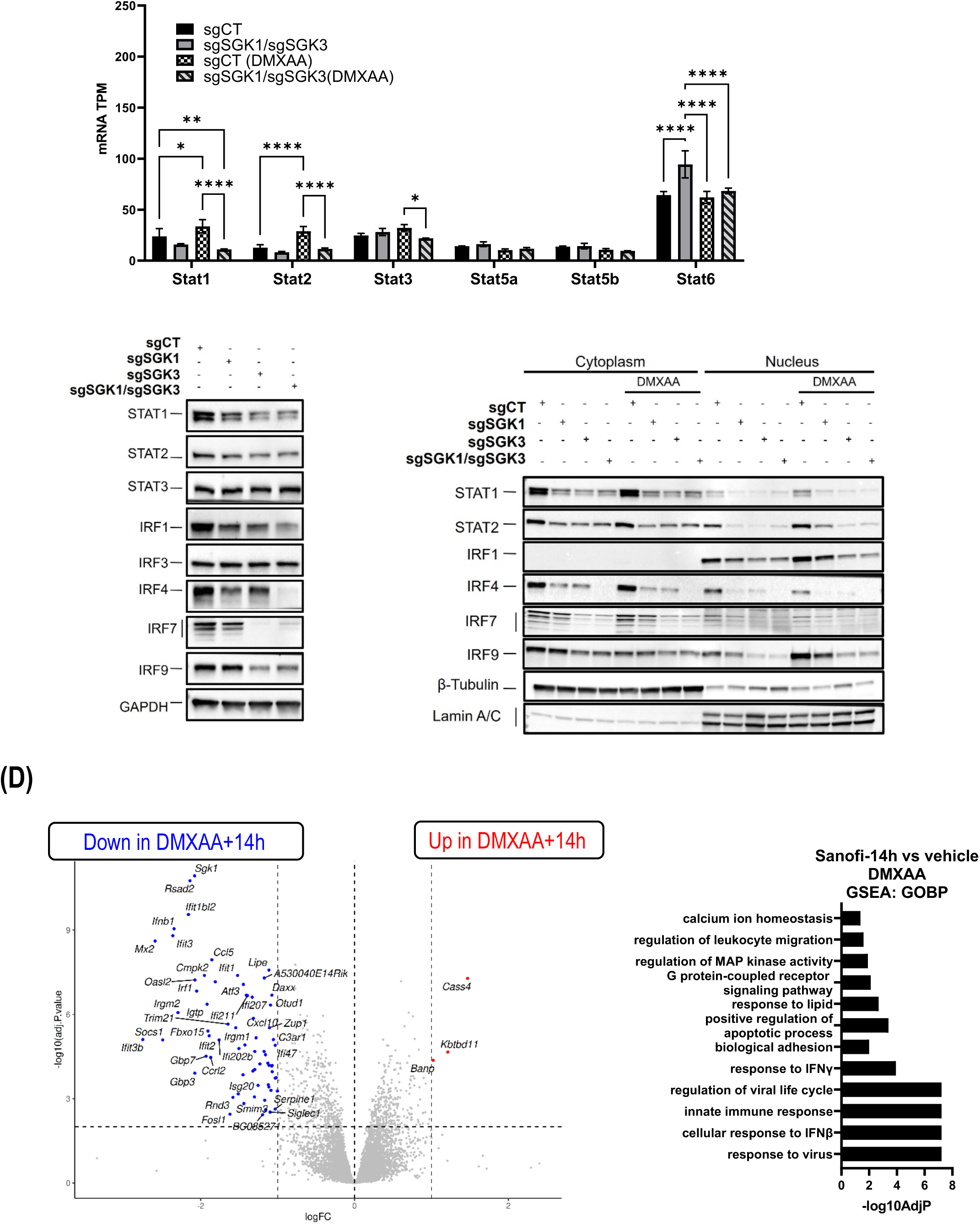
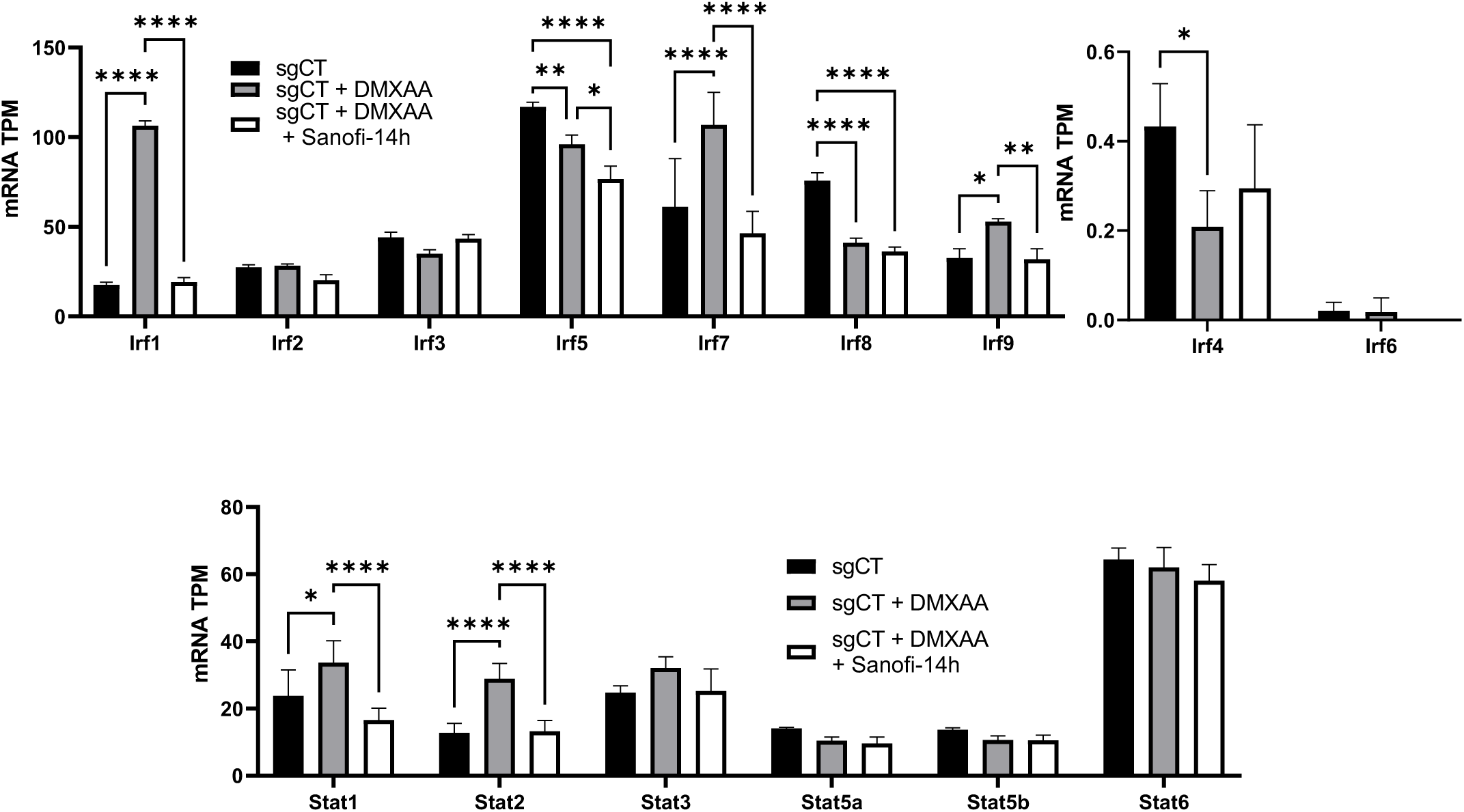
SGK1 and SGK3 KO blocks expression of Interferon Stimulated Genes (ISGs) and innate immunity regulators. **A)** Heatmap (left panel) reflecting the differentially expressed genes after knocking out SGK1/3 without any stimulation. Blue indicates genes that are being downregulated while red show genes that are upregulated in the dKO cells. Volcano plot (middle graph) depicting the expression of Interferon Stimulated Genes (ISGs) at basal levels in SGK1/3 dKO cells plotted by log fold change(logFC) in the x-axis and -log of adjusted p value in the y-axis. Right graph depicts the Gene Set Enrichment Analysis (GSES) and Gene Ontology Biological Process (GOBP) representing the most changes pathways after SGK1/3 dKO in non-stimulated cells based on -log10 of the adjusted p value in the x-axis. **B)** Volcano plot (left graph) showing the expression of ISGs after DMXAA treatment for 2 h in SGK1/3 dKO cells. Significance was achieved with an adjusted p value of <0.05 and a fold change >2 in either direction. GSES and GOBP analysis (right graph) of the most affected pathways after DMXAA treatment in SGK1/3 dKO cells. **C)** IRFs and STATs mRNA expression in Transcripts Per Million (TPM) (*p<0.05) in SGK1/3 dKO cells with and without DMXAA(2h). Immunoblot of IRFs and STATs protein expression at basal conditions (left panel) and their nuclear localization (right panel) after DMXAA treatment for 1h. **D)** Volcano plot (left graph) showing genes that are upregulated (red) and downregulated (blue) by Sanofi-14h on the DMXAA response in RAW264.7 cells. GSES and GOBP analysis (right graph) reflecting the effects of Sanoni-14h on the most significant DMXAA responsive pathways. IRFs and STATs mRNA-TPM expression with and without Sanofi-14h after DMXAA treatment. All experiments were repeated at least three times.

We next examined gene expression by RNAseq in STING-activated cells and found striking loss of IFN and ISG pathway genes in SGK1/3 dKO cells stimulated with DMXAA relative to WT cells [**Figure 3B**]. The extent and significance of downregulation is enhanced after DMXAA stimulation relative to the basal state. GSEA confirmed the loss of interferon and host defense pathway transcription in SGK1/3 dKO cells. Other gene sets impacted by SGK1/3 deletion include those involved in antigen processing, T cell stimulation, cell adhesion, and cell differentiation. Examination of individual transcription factors showed that in addition to the baseline defect in expression of Irf1, Irf7, Irf9, Stat2, and Stat6, SGK1/3 dKO macrophages do not upregulate Irf7 or Stat2 RNA after DMXAA treatment [**Figure 3B and 3C**]. We extended these findings by immunoblotting which demonstrated a baseline decrease of STAT1, STAT2, IRF1, IRF7, and IRF9 protein levels in dKO cells [**Figure 3C**]. SGK3 appears to control IRF7, IRF9 and STAT2 preferentially as compared with SGK1. Despite relatively low SGK1 protein expression in RAW264.7 cells, SGK1 KO cells show enhanced loss of IRF4 as compared with SGK3 KO cells. Consistent with the expression defects, we observed that SGK1/3 deletion diminished nuclear translocation of IRF4, IRF9, STAT1 and STAT2 and IRF1 in SGK1/3 KO cells [**Figure 3C**]. IRF9 and STAT1 are part of the Interferon Stimulated Gene Factor 3 (ISGF3)^42^, which is responsible for inducing expression of Interferon Stimulated Genes (ISGs), IRF1 is also involved in the regulation of ISGs in an IFN dependent and independent manner^43^. STAT2 is also part of the ISGF3 complex^42^ which partially explains (along with a block in IRF1 expression) why the vast majority of affected genes were ISGs. Together, these results indicate that SGK1 and SGK3 regulate the expression of IFNβ and ISGs by controlling expression of innate immune regulators IRF1, IRF4, IRF7, IRF9, STAT1 and STAT2 at the RNA level and prior to STING-induced. Additional immune-related mechanisms could depend on SGK1 and SGK3, both basally and under induced conditions, in macrophages.

### Sanofi-14h blocks DMXAA-stimulated Interferon and ISG gene expression in macrophages

We next asked whether effects of Sanofi-14h treatment overlap the observed transcriptomic effect of SGK1/3 deletion in RAW264.7 cells. Because high dose (5-10μM) Sanofi-14h efficiently blocks TBK1 activation [**Figure 1C**], we selected a lower dose (500nM) of Sanofi-14h that inhibits SGK1 and SGK3 (as assayed by NDRG1 phosphorylation) but largely leaves TBK1 activation intact. Similar to SGK1/3 deletion, 500nM Sanofi-14h inhibited the inducible expression of Ifnβ1 and Stat1 as well as several ISGs (including Mx1, Gbp5, ISG15, Ifit1/2/3, Rsad2) and chemotaxis-related genes (Ccl5, Cccrl2) [**Figure 3D**]. GSEA analysis confirmed the similar transcriptomic effect of SGK inhibition and deletion, showing that the most affected genes are those related to pathogen sensing. Moreover, RNAseq confirmed that Sanofi-14h at a dose of 500nM has a similar effect as deletion of SGK1 and SGK3 in terms of IFN and ISG expression. However, only Irf1, Stat1 and Stat2 RNA are impaired by Sanofi-14h to a similar level as in SGK dKO cells. Irf7 and Irf9 RNA levels were not significantly reduced by Sanofi-14h during the short (2 h pretreatment, 2 h stimulation) exposure to drug, suggesting they are not wholly responsible for Ifnβ1 regulation by Sanofi-14h in this timeframe[**Figure 3D**]. The less broad effect of Sanofi-14h is likely explained by acute responses of the cells versus the stable loss of SGK1/3. These data confirm that Sanofi-14h and SGK1/3 regulate STING-driven gene expression responses in macrophage cells.

### Sanofi-14h disrupts STING-stimulated TBK1-STING interaction

In the STING-driven antiviral immune response TBK1 is induced to interact with STING to undergo trans-autophosphorylation, to phosphorylate STING and to activate IRF3^39^. We asked whether doses of Sanofi-14h that inhibit TBK1 activation do so by blocking induced STING-TBK1 interaction. To address this, we pre-treated RAW cells with 5μM Sanofi-14h prior to stimulation with DMXAA followed by co-immunoprecipitation. DMXAA induced robust interaction between STING and TBK1 as expected, and Sanofi-14h efficiently impaired DMXAA induced TBK1-STING interaction [**Figure 4A**]. Additionally, Sanofi-14h blocked DMXAA phosphorylation of STING as indicated by a reduction in the slower migrating STING form [**Figure 4A**]^42^. We next examined whether Sanofi-14h treatment alters subcellular trafficking of TBK1 and IRF3 following STING activation. We isolated cytoplasmic and membranous (containing the ER, mitochondria and Golgi) fractions after DMXAA treatment. We observed recruitment and phosphorylation of TBK1 and IRF3 in the membrane fraction as expected [**Figure 4B**]. Pretreatment with Sanofi-14h prior to DMXAA prevented TBK1 and IRF3 translocation/retention and subsequent STING activation in the ER [**Figure 4B**]. These outcomes support the hypothesis that Sanofi-14h inhibits STING pathway signaling by preventing TBK1-STING interaction in the ER, thus blocking subsequent recruitment and phosphorylation of IRF3 and inhibiting this key step in the activation pathway.

**Figure 4:**
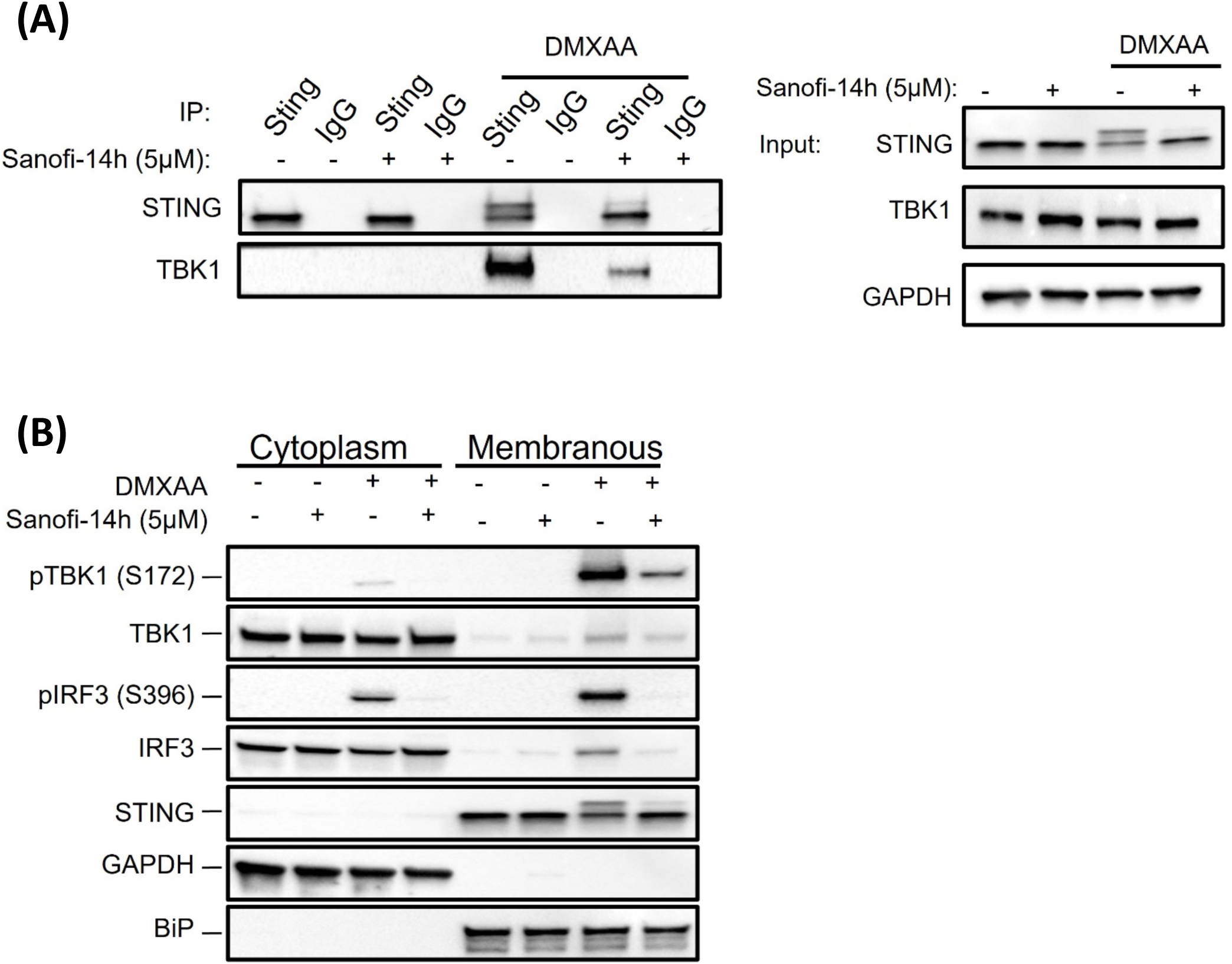
Sanofi-14h disrupts TBK1/STING interaction. **A)** Effects of Sanofi-14h treatment on STING and TBK1 interaction of DMXAA treatment for 1h. STING was immunoprecipitated and the interaction was measured based on TBK1 detection in immunoblot. IgG was used a negative control and inputs show the levels of both proteins in total lysate. **B)** RAW264.7 cells were fractionated and immunoblotted for TBK1 and IRF3 translocation into the ER-Golgi Space after Sanofi-14h(2h) and DMXAA treatment for 1h.

### Class effects of SGK inhibition on STING/TBK1/IFN signaling

In order to address mechanisms associated with the ability of Sanofi-14h to block the STING/TBK1 activation step downstream of DMXAA treatment, we analyzed other inhibitors of AGC family kinases. Using an IRF3-dependent luciferase reporter system in RAW264.7 macrophages (Invivogen), we tested DMXAA-stimulated cells pre-treated with 5 different SGK inhibitors across a 10,000-fold concentration range. For this study we synthesized and tested Sanofi-17a, a structural derivative of Sanofi-14h, that was recently described^31^ and which preferentially targets SGK1 over SGK3. We included three other small molecules, namely GSK650394, SI113, and EMD638683, with established SGK inhibitory activity^27–29^. We found that each of these known SGK inhibitors, with the exception of EMD638683, blocked DMXAA-stimulated IRF3-promoter activity with an IC_50_ of 3.334 μM and 5.612 μM, respectively [**Figure 5A**]. Sanofi-17a was less effective than the other SGK inhibitors in the reporter assay with an IC_50_ of 9.006 μM whereas Sanofi-14h was the most effective with an IC_50_ of 1.728 μM. Since Sanofi-17a is a derivative of Sanofi-14h, we examined the effects of Sanofi-17a on STING-activated signaling. Similar to Sanofi-14h, Sanofi-17a at low micromolar concentrations effectively blocked NDRG1 phosphorylation [**Figure 5B**]. Sanofi-17a inhibited TBK1, STAT3, and IRF3 phosphorylation induced by DMXAA with 50% inhibition of TBK1 phosphorylation seen at ∼10μM, similar to but less effective as compared with Sanofi-14h [**Figure 5B and 1C**].

**Figure 5:**
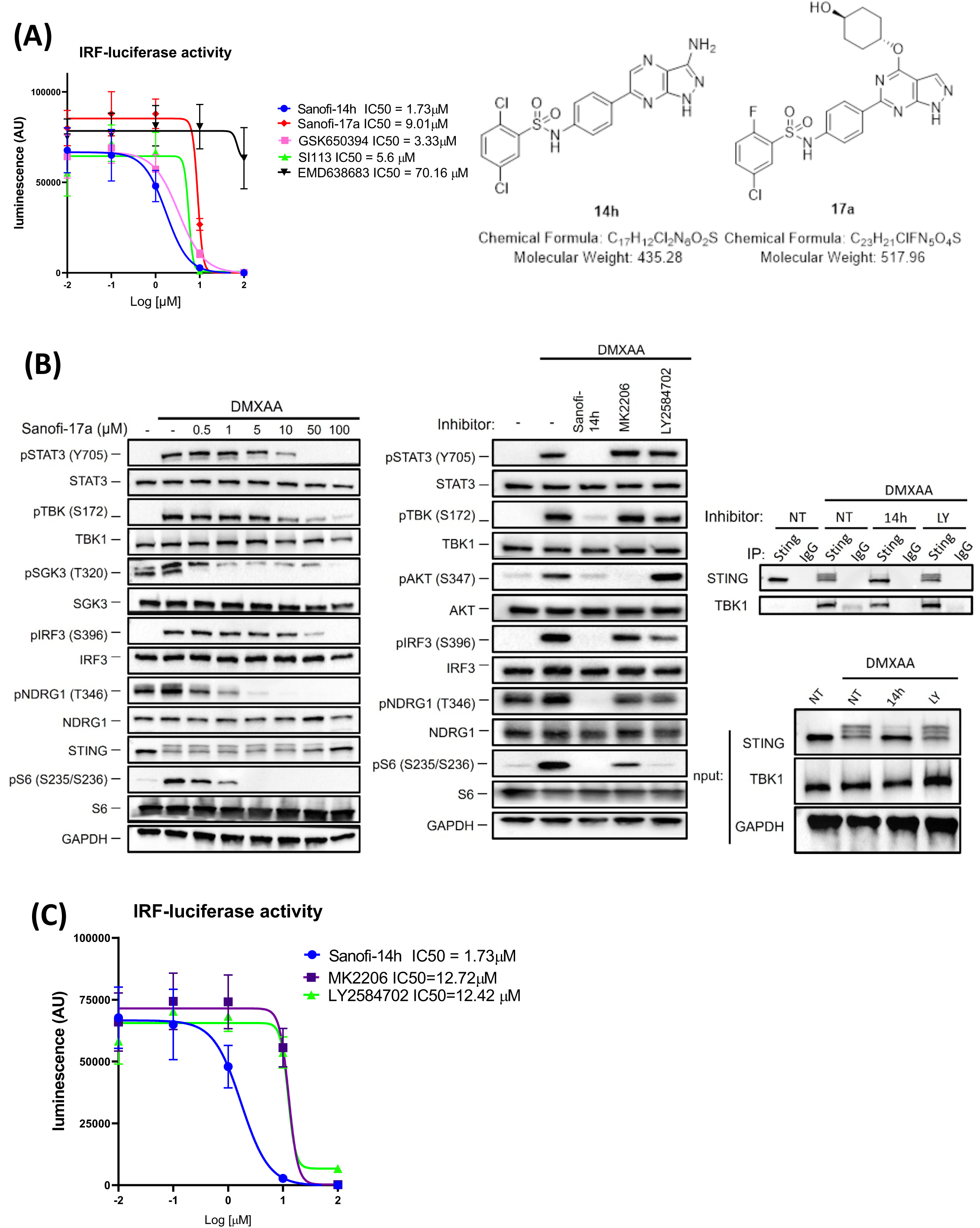
Other SGK inhibitors have variable effect on IRF DMXAA driven activation. **A)** IRF activation luciferase assay (left graph) in the presence of diverse SGK inhibitors and DMXAA stimulation. Raw-Dual Luciferase cells were seeded in 96 well plates, left to rest overnight and treated with the different inhibitors at different doses for 2 h and then treated with DMXAA for 24h. Luciferase signal directly correlates with type I production and the graph depict Log10 of the drug concentrations. Chemical Structure of Sanofi-14h and closely related compound Sanofi-17a (right panel). **B)** Immunoblots (left panel) showing the effects of Sanofi-17a, LY258702 and MK2206 on markers after DMXAA treatment. Cells were treated with different doses of Sanofi-17a and with 10 μM of Sanofi-14h, MK2206 and LY2584702 for 2 h and the treated with DMXAA for two additional hours. Co-immunoprecipitation (right panel) showing the effects of LY258702(2 h pre-treatment) on TBK1/STING interaction after DMXAA treatment for 1 h. **C)** Dual Luciferase assay depicting the effects of the same inhibitors on IFNβ production. The graph is depicted the same way as **A.**

It was reported that interaction between the TBK1/STING complex with IRF3 is dependent on the structural activity of P70S6K1^44^ without affecting TBK1/STING interaction. In this regard, we examined phosphorylation of ribosomal S6 protein, a substrate of the S6 kinase family, based on reports that P70S6K regulates the STING/TBK1/IRF3 complex^44^. DMXAA stimulated S6 phosphorylation in RAW264.7 cells and this was blocked by Sanofi-17a in the low micromolar range [**Figure 5B**]. Sanofi-17a also impaired DMXAA-stimulated phosphorylation of SGK3 T320 at 500nM [**Figure 5B**]. SGK3 T320 is reported to be a direct phosphorylation substrate of 3-phosphoinositide-dependent protein kinase 1 (PDK1)^45^. The sensitivity of SGK3 T320 phosphorylation to inhibition by Sanofi-17a may represent either loss of feedback from SGK3 to PDK1 or an undescribed component of SGK3 autophosphorylation at that site. Thus, Sanofi-17a blocks phosphorylation of S6 around 1μM but is less effective at blocking phosphorylation of TBK1 and IRF3.

Because SGK inhibitors are reported to have activity against other AGC kinases like P70S6K1 or Akt^23^, we asked whether the signaling effects of Sanofi-14h treatment overlap with those of Akt or P70S6K inhibitors. In RAW264.7 cells stimulated with DMXAA, Akt inhibition by MK2206 or P70S6K inhibition by LY2584702 had no effect on STAT3 Y705 phosphorylation while Sanofi-14h at the same dose completely blocked phospho-STAT3 [**Figure 5B**], consistent with loss of IFNβ production. Akt inhibition had no measurable effect on DMXAA-stimulated phosphorylation of TBK1 or IRF3. P70S6K inhibition partly abrogated IRF3 phosphorylation while leaving TBK1 phosphorylation largely intact, potentially consistent with reports that S6K1 interacts with STING^44^. When assayed using the IRF3-luciferase reporter, Akt and P70S6K inhibitors had weak effects on reporter activity with IC_50_ greater than 50μM, while Sanofi-14h was significantly more potent [**Figure 5B**]. To further test whether P70S6K inhibition has the same effect as Sanofi-14h we assayed the interaction between TBK1 and STING by co-immunoprecipitation. LY2584702 did not alter the STING-TBK1 interaction after DMXAA treatment [**Figure 5C**], suggesting that Sanofi-14h is unlikely to work by the same mechanism as S6K inhibition. The lack of effect on TBK1 phosphorylation, IFN expression, or STING-TBK1 interaction by Akt or P70S6K inhibitors makes it unlikely that Sanofi-14h affects the TBK1/IFN pathway via those kinases (see Discussion).

## DISCUSSION

Inhibitors of the SGK family have been developed to target these kinases in cancer as well as other disease indications, such as osteoarthritis. As examples, Bago et al^23^ used Sanofi-14h to block induced SGK activity associated with Akt inhibition in breast cancer, and Halland et al ^23, 31^ developed Sanofi-17a to target SGK in an osteoarthritis model. Sanofi-14h is preferential for SGK3 and Sanofi-17a is preferential for SGK1, underscoring the potential importance of SGK subtype preference for these inhibitors related to different biological effects and expression. These studies showed that other kinases are targeted by these inhibitors, as Sanofi-14h was shown to have significant activity against S6K1 and MLK3 ^23, 31^. Our studies revealed an unexpected effect of Sanofi-14h on inhibition of IFNβ mRNA induction following treatment of macrophages cells with the STING agonist DMXAA (Fig. 1A) and this result was extended to corresponding effects on viral replication and virus-induced IFNβ (Fig. 1B). Sanofi-14h clearly targeted SGK in the macrophage cells as shown through its ability to block phosphorylation of the SGK target NDRG1.

To address a mechanism for the inhibition of IFNβ mRNA induction by Sanofi-14h, we analyzed the effects of Sanofi-14h on DMXAA-induced signaling and found this compound blocked phosphorylation of TBK1 and IRF3 as well an IFN-driven STAT3 phosphorylation (Fig. 1C). Notably the effect of blocking of pNDRG1 was nearly complete at a 500nM dose while the inhibitory effects on TBK1 signaling were only partial at that dose, suggesting a different target in controlling TBK1 activation. The effect of Sanofi-14h was shown to be at the level of blocking the induced interaction between STING and TBK1 (Fig. 4), thus at an early stage in the STING-induced signaling pathway (and see below).

Analysis of RAW264.7 cells revealed that they express SGK3 at relatively high levels and SGK1 at significantly lower levels. SGK2 expression (RNA and protein) was not detectable in these cells. Interestingly, SGK1 mRNA and protein are induced in the DMXAA-response (Fig.1C and sup 1A), suggesting an important role for SGK1 in the innate immune response as an early response gene. To gain insight into potential mechanisms associated with SGK1/3 in the STING-induced pathway, we used CRISPR-CAS9 to knock out expression of these genes in RAW264.7 cells. Consistent with the Sanofi-14h response, loss of SGK1/3 blocked STING-induced IFNβ mRNA induction (Fig. 2A). Yet, loss of SGK1/3 in the macrophage cells did not block the induction of phosphorylation of TBK1 and IRF3, indicating an SGK-independent mechanism for this key regulatory event (Fig. 2D).

To gain insight into regulatory roles for SGK1/3 in the STING-induced response, we performed RNA sequencing on the dKO cells at baseline and under DMXAA-stimulated conditions and compared the results to WT cells (Fig. 3). In SGK1/3 KO we found low expression of direct and indirect regulators of IFNβ production IRF1, IRF4, IRF7, IRF9, STAT1 and STAT2 at basal and stimulated conditions at the protein and RNA levels. Among them, IRF7 is essential for the production of IFNβ mRNA in response to STING activation because it directly engages the IFNβ promoter along with IRF3^46^. Like IRF7, other IRFs and STATs were more dependent on SGK3 as shown by their decrease in protein levels in the SGK3 KO cells, whereas IRF4 appears dependent on SGK1. Thus, results where loss of SGK1 expression leads to loss of IFNβ expression (Fig. 2B) may be related to the corresponding loss of expression of IRF4, a transcription factor involved in macrophage polarization^47^. IRF1 is essential for the expression of ISGs such as OAS2, BST2 and RNASEL as well as IRF9^43^ which along with STAT1 and STAT3 forms a complex termed ISGF3 responsible for IRF7 expression^48^ . These results indicate that SGK1 and SGK3 are involved in establishing the expression of a variety of key transcription factors both constitutively and inducible in macrophage cells, thus coordinating the innate immune response.

To address whether SGK1/3 play a role in immediate STING-induced signaling, we treated cells with 500nM Sanofi-14h, a dose that is preferential for targeting SGK1/3 over blocking TBK1 activation, and performed RNAseq studies. The results demonstrated that Sanofi-14h blocked expression of a subset of genes within the larger group affected by SGK1 and SGK3 knockout such as Irf1, STAT1 and STAT2, which were inhibited prior to STING activation in dKO cells. In comparison, a short pre-treatment with Sanofi-14h did not impair IRF7 or IRF9 expression, though these transcripts are under-expressed in SGK KO cells (Fig. 3). This may relate to the short time frame of inhibition compared to CRISPR deletion and to the half-life of Irf7 and Irf9 mRNAs. Likewise, the effects of Sanofi-14h at 500nM were phenotypically similar to those of the dKO cells in terms of the gene sets that were affected after DMXAA treatment. IFNβ production, response to viral cycle, and response to bacterium were among such gene sets. These data suggest that SGK controls expression of IFNβ mRNA by positively regulating IRF7 expression and this may occur via a mechanism involving IRF1, IRF9, STAT1 and STAT2 but future experiments will be necessary to define this response.

The ability of Sanofi-14h treatment to inhibit activation of TBK1 led us to examine a panel of SGK and AGC family kinase inhibitors for their effects on STING-induced signaling. In this regard, we synthesized Sanofi 17a^31^, a derivative of Sanofi-14h, which preferentially targets SGK1 (Fig. 5A)^31^. We found that Sanofi-14h, Sanofi-17a, GSK650394, and SI113 all inhibited STING-driven IRF3 activation in a reporter assay with similar potency. The SGK inhibitor EMD638683, reported to exhibit preference for SGK1 over SGK3 ^49^, was ineffective at blocking IRF3 activation (Fig. 5A). Sanofi-17a blocked phosphorylation of TBK1 and IRF3, although seemingly less efficiently than Sanofi-14h, with inhibitory effects around 10 μM (Fig. 5B) whereas Sanofi-14h blocks TBK1 and IRF3 in the ∼1 μM range (Fig. 1A). Consistently, Sanofi-17a blocked the IRF3-dependent reporter less effectively than did Sanofi-14h (Figure 5A). Overall, our studies show that structurally dissimilar SGK inhibitors such as Sanofi-14h, SI113, and GSK650394 all inhibit the STING-induced innate immune response.

It was reported that the AGC family kinase S6K1 associates with STING to promote recruitment of IRF3, but not TBK1, into the STING complex, in the viral-regulated innate immune response^44^. Interestingly, this mechanism was reported to not require the catalytic activity of S6K1. In this regard, we analyzed effects of different AGC kinase inhibitors on phosphorylation of S6 and on the DMXAA-induced downstream signaling response. Sanofi-14h has been shown (in cell-free assays) to block S6K1 in a range similar to that of SGK3 and SGK1^23^, while Sanofi 17a was reported to have little to no effect on S6K1 activity ^31^. Interestingly, we observed that phosphorylation of S6 is induced by DMXAA (Fig. 5B) suggesting the involvement of an S6 kinase in the STING-induced response. Both Sanofi-17a and Sanofi-14h inhibited the phosphorylation of S6 induced by DMXAA. The Akt inhibitor MK2206 prevented phosphorylation of S6 (though not as effectively as Sanofi-14h) but did not block the induced phosphorylation of TBK1 or IRF3 in the STING pathway. The P70S6K inhibitor LY2584702 effectively impeded the induced phosphorylation of S6, but less effectively blocked phosphorylation of IRF3. Consistently neither the Akt inhibitor nor the P70S6K inhibitor affected phosphorylation of STAT3 or activation of the IRF3 reporter. While Sanofi-14h impaired recruitment of TBK1 to the STING complex in response to STING stimulation, LY2584702 did not inhibit recruitment of TBK1 and did not hinder phosphorylation of STING which is controlled by TBK1 recruitment (Fig. 5B). These studies show that S6K and Akt inhibition does not phenocopy SGK inhibition and suggests that Akt and S6K do not mediate the effect of Sanofi-14h and related compounds on TBK1 and IRF3 activation. Our study demonstrates SGK1 and SGK3 as important, targetable regulators of the STING/IFN pathway. Future studies are directed at the mechanism of action of Sanofi-14h, and other SGK/AGC inhibitors, relative to inhibiting TBK1 and IRF3 activation downstream of activated STING.

We propose a model in which SGK1/3 control the expression of key regulators of the cGAS/STING pathway (Figure 6). Upon viral infection cGAS recognizes dsDNA and produces 2’3’-cGAMP which activates STING, TBK1 then gets recruited to STING in the ER where it phosphorylates IRF3 which subsequently dimerizes and translocates to the nucleus to initiate type I IFNs transcription. TBK1 also phosphorylates IRF7 through an unknown mechanism. Sanofi-14h blocks activation of TBK1 and consequent IRF3 and IRF7 phosphorylation by interacting with a regulator other than SGK1/3. IRF7 protein levels, however, are maintained by SGK1/3 at basal conditions. Either the use of Sanofi-14h or the genetic ablation of SGK1/3 renders the cells susceptible to viral infection. This works implicates currently existing SGK inhibitors as useful modulators of IFN signaling and points to the SGK kinase family as a useful target in IFN-related diseases.

**Figure 6:**
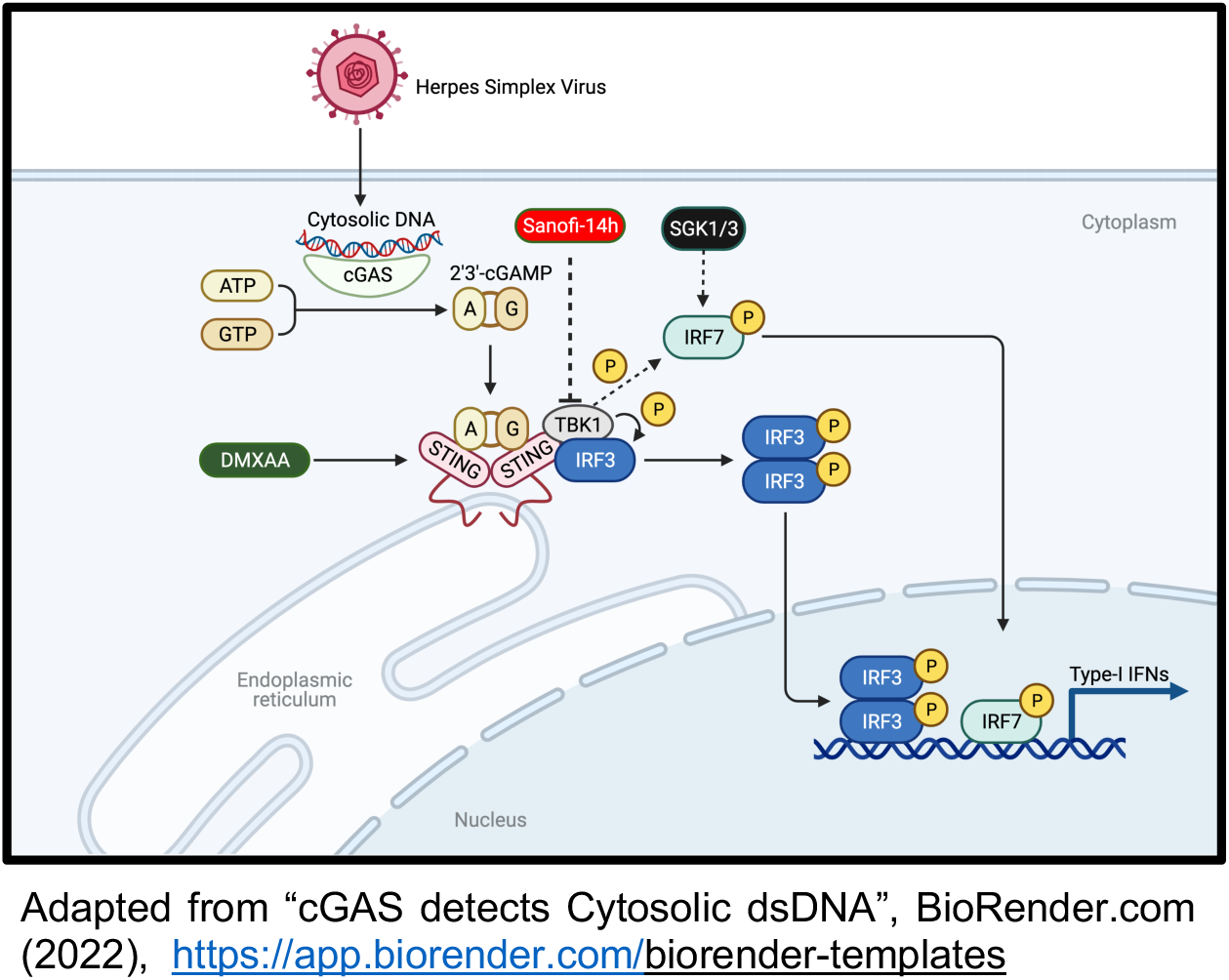
Model of SGK regulation of STING/TBK1 signaling. Sanofi-14h inhibits phosphorylation of TBK1 through an unknown mechanism preventing STING and IRF3 phosphorylation, dimerization and nuclear translocation in the presence of 2’3’-cGAMP and DMXAA. SGK1 and SGK3 promote IRF7 expression and stabilization under basal and DMXAA stimulated conditions in macrophages.

## Materials and Methods

### Cell culture

RAW 264.7 (passage 13), Vero and HEK293T/17(passage 30) cells were purchased from the tissue culture facility at UNC. All cell lines were cultured in DMEM High glucose medium (Gibco, 11995-065) supplemented with 10% fetal bovine Serum (FBS) (Avantor, 97068-085) and 1% penicillin/streptomycin(P/S) (Gibco, 15140122) (regular DMEM) in an atmosphere of 5% CO_2_ and 95% humidity at 37°C. RAW 264.7 cells were maintained in 10 cm plates and passaged with a scraper after reaching 90% confluency from passage 13 until passage 35. DMXAA (Cayman Chemical, 14617) treatments in RAW 264.7 cells were carried out at a 10 μg/mL dose for all experiments at a density of 3.5x10^6^ cells in 60 mm plates (immunoblots, RT-qPCR and RNA seq) and 35x10^6^ cells in 10 cm plates (co-IP and cell fractionations). Treatments with Sanofi-14h (MRC Pure Reagents) were performed at the indicated doses. HEK293T/17 cells were maintained in 10 cm plates and passaged after reaching 90% confluency with 2 mL of Trypsin-EDTA (0.25%) (Gibco, 25200056) at 37°C for 2 minutes until passage 50. Cells were regularly tested for mycoplasma. RAW-Dual Lucia (Invivogen, rawd-ismip) cells were maintained in DMEM complemented with 100 μg/mL of Normocin (Invivogen, ant-nr-1), 10% heat inactivated FBS, 200μg/mL of Zeocin (Invivogen, ant-zn-05) and 1% penicillin/streptomycin(P/S). For the luciferase assays RAW-Dual Lucia cells were plated in 96 well plates at a density of 0.2x10^6^ cells per well and let to rest until the next morning. Cells were treated with the indicated inhibitors for 2 hours and them stimulated with DMXAA for 24h. The luciferase assay was performed following the manufacturer’s instructions.

### Bone marrow derived macrophages preparation

The SGK3 knockout mice were a kind gift from Dr. David Pearce at the University of California San Francisco and have been described previously^50^. All procedures involving mice handling were in accordance to approved protocols by the University of North Carolina Chapel Hill (IACUC protocol 20-167). Bone marrow from WT and SGK3 KO mice was extracted as previously described ^51^. Bone marrow was harvested from mice as previously described^52^, and cultured in L929-cell condition media (LCCM) to generate macrophages ^53^. Cells were seeded in non-tissue culture treated 10 cm plates (Fisherbrand, FB08755712) in media (complete media) containing RPMI1640(Corning, 10-040-CV), 20% LCCM, 10% FBS, 1%(P/S), 2 mM of L-Glutamine (Gibco, 2503-081), and 55 μM of 2-Mercaptoethanol (Gibco, 21085023). After 24 hours cells were washed twice with PBS to remove non-adherent cells and fresh complete media was added. Media was changed on days 3 and 5 prior to scraping, counting, and replating. On day 7 media was changed to regular media without LCCM for 24 hours prior to experiments.

### Cloning

Guide RNAs targeting mSGK1 exon 5(GATTTCAGGTTCTTCTGGCT) and mSGK3 exon 3 (TTCAAGACATTAAATGCAG) were designed using the online tool CHOPCHOP (http://chopchop.cbu.uib.no/) and cloned into TLCV2 (Addgene plasmid # 87360,) as described previously^5455^. Plasmids were transformed into One Shot™ Stbl3™ Chemically Competent *E. coli* (Invitrogen, C737303) and plated in ampicillin(100ug/mL) plates overnight at 37°C. Colonies were picked and sequenced using the LNX primer (AGCTCGTTTAGTGAACCGTCAGATC).

### Lentivirus production

Lentiviral particles were produced as described with some modification^56^. HEK293T/17 cells (0.75x10^6^ per well) were plated in 6 well plates and after 20 hr cells were co-transfected with the 2μg of TLVC2-gRNA, 1μg of psPAX2 (Addgene #12260) and 0.5μg pMD2.G (Addgene #12259) in a ratio of 4:2:1 using FuGENE HD and OptiMEM according to the manufacturer’s instructions. Media was changed the next day and after 48h the viral particles were collected by filtration (0.45 μM filter).

### CRISPR KO cell line generation

RAW 264.7 cells (passage 13) were plated at a density of 0.75x10^6^ cells per well in a 6 well plates and were infected using 1 mL of lentivirus, 1 mL of fresh regular DMEM, 8μg/mL Polybrene (Santa Cruz Biotechnology, sc-134220) and spin infection (1372 RCF for 1hr at RT). Cells were re-infected the next day and selected the following day with Puromycin(2.5μg/mL). After the non-infected control died a portion of the cells was frozen with BAMBANKER FREEZE freezing media (Fisher Scientific, NC9582225) and the rest was plated in 10 cm without puromycin and supplemented with 1μg/mL of doxycycline which was refreshed every day for 7 days. Cells were sorted based their GFP “high” signal and plated in 96 well plates at a ratio of 1 cell/well. Clones were screened for SGK1 or SGK3 KO through immunoblot (Supplemental figure). The double SGK1 and SGK3 KO RAW 264.7 cells were generated by Synthego Corporation (Redwood City, CA, USA) as previously described^57^ and single clones were isolated in the same way as the one for the single KOs.

### Immunoblot

Immunoblots were performed utilizing lysis buffer (50 mM Tris-HCl pH 7.4, 150 mM NaCl, 1mM EDTA, 1mM EGTA, 1 mM Beta-Glycerol-Phosphate, 2.5 mM Sodium-Pyrophosphate, 1% Triton-100X, 0.1% SDS) complemented with fresh protease (Sigma, 11697498001) and phosphatase inhibitors (Sigma, P0044-5ML). Proteins were transferred using the Trans-Blot Turbo RTA Mini 0.2 µm Nitrocellulose Transfer Kit (Bio-Rad, 1704270) in the Trans-Blot Turbo Transfer System (Bio-Rad 1704150). Membranes were then blocked for 1 hat RT in 5% dry milk and blotted with primaries overnight at 4 °C, after three washes with TBST the membranes were blotted with secondary for 1 hat RT in blocking buffer. The three washes were repeated and the membranes were imaged using Clarity Western ECL (Bio-Rad, 1705060) in the ChemiDoc Imaging System. Images were processed with the ImageJ software (https://imagej.nih.gov/ij/). The following primary antibodies were from Cell Signaling Technologies and were used 1:1000 unless otherwise indicated: phospho-STAT3 Thr705 (9145S), STAT3 (12640S), phospho-TBK1 Ser172 (5483S), TBK1 (38066S), phospho-IRF3 Ser396 (29047S), IRF3 (4302S), STING (50494S), phosphor-NDRG1 Thr346(5482), NDRG1(9408, 1:500), GAPDH (5174S), β-Tubulin(2146S), Lamin A/C (4777), SGK1 (12103), SGK3 (8154), BiP(3177T), STAT1 (14994S), STAT2 (4597S), IRF1 (8478S), IRF4(62834T), IRF7 (72073) and IRF9 (28845S). Anti-Rabbit (W4018) and Anti-Mouse (W4028) secondaries antibodies were bought from Promega.

### Co-immunoprecipitation

STING immunoprecipitation was performed as previously described ^44^. In brief, after treatment samples were lysed in IP buffer (20 mM Tris-HCl pH 7.4, 137 mM NaCl, 1 mM EDTA, 1% NP-40, 10% glycerol) complemented with fresh protease and phosphatase inhibitors. 700 μg of protein were mixed and rotated with 0.5 μg of STING antibody for 3 h at 4 °C after which 25 μL of Dynabeads Protein G (Invitrogen, 10004D) were added to each sample and rotated for an additional hour. Samples were washed three times for 5 min with IP buffer at 4 °C. After the last wash, the supernatant was discarded and samples were boiled in 4x loading buffer with 10% of 2-Mercaptoethanol for 5 min prior to SDS-PAGE and immunoblotting. Clean-Blot™ IP Detection Reagent (Thermo Fisher, 21230) was used for detection to avoid Ig heavy chain.

### Nuclear fractionation

Cytoplasmic and nuclear fractions were obtain as previously described^58^ with modifications. Cells were washed with PBS and centrifuged at 2,300 RCF for 30 seconds and the supernatant discarded. Cells were resuspended in 250 μL of 0.1% NP-40 diluted in cold PBS (with protease and phosphatase inhibitors) and centrifuged at 4 °C at 2000 RCF for 2 min. The supernatant (cytoplasmic fraction) was removed without disturbing the nuclear pellet. The nuclear pellet was washed by gently resuspending with 1 mL of the 0.1% NP40 buffer and samples were centrifuged for 2 min at 2,000 RCF at 4 °C. The supernatant was discarded and the pellets were resuspended in RIPA buffer and sonicated for 10 min (at high with 30 sec on and 30 Sec off) in a water bath sonicator. After sonication pellets were centrifuged 4 °C at 16, 000 RCF for 10 minutes and protein from both fractions were quantified and immunoblotted. β-Tubulin was used as a cytoplasmic control while Lamin A/C was used as nuclear control.

### ER fractionation

Cytoplasmic and membranous fractionation was performed as previously described^59^. Cells were washed with PBS after treatment, collected and centrifuged at 2, 300 RCF for 30 seconds. The supernatant was removed and cells were resuspended in buffer 1 (200μg/mL Digitonin, 150 mM NaCl, 50 mM HEPES pH7.4 (Boston Bioproducts, NC0703855), ddH2O, fresh protease and phosphatase inhibitor) and put on ice for 10 min. Afterwards, they were centrifuged for 2 min at 2000 RCF at 4 °C and the supernatant (cytoplasmic fraction) was removed and the pellets were washed with 1 mL of cold PBS. Pellets were resuspended in buffer 2 (1% NP-40, 150 mM NaCl, 50 mM HEPES pH7.4, ddH2O, fresh protease and phosphatase inhibitor) and left on ice for 30 min. After centrifugation, the supernatant was kept (membranous organelles) and the pellets were discarded. Samples were then quantified and immunoblotted.

### Dimerization

The IRF3 dimerization procedure was adapted from the literature^60^ as follows: after treatment cells collected and lysed in dimerization Buffer (50 mM Tris-HCl pH 7.4, 150 mM NaCl, 1mM EDTA, 1mM EGTA, 1 mM Beta-Glycerol-Phosphate, 2.5 mM Sodium-Pyrophosphate, 1% Triton-100X) for 10 min on ice. Samples were centrifuged for 10 min at maximum speed at 4 °C and protein was quantified. Proteins were mixed with 2x native sample buffer (Bio-Rad, 1610738) and resolved for 35 min at 200 V in precast gels with native running buffer (25 mM Tris and 192 mM glycine) complemented with 0.2% sodium deoxycholate in the cathode chamber prior to transfer and immunoblotting.

### Reverse Transcriptase Quantitative PCR (RT-qPCR)

Total RNA was extracted using the Quick-RNA miniprep kit (Zymo Research, R1055). cDNA was prepared with the iScript™ cDNA Synthesis Kit (Bio-Rad, 1708891) and RT-qPCR was performed utilizing the iTaq™ Universal Probes Supermix (Bio-Rad, 1725131) according to the manufacture’s instruction. Mouse Taqman probes were ordered from Thermo Fisher: IFNβ (Mm00439552_s1), SGK1 (Mm00441380_m1) and GAPDH (Mm99999915_g1).

### Herpes Simplex Virus (HSV)

HSV-1 KOS ^61^ was a kind gift from Dr. Steve Bachenheimer (University Of North Carolina at Chapel Hill) and was expanded as described^62^. Briefly, Vero cells were grown to 100% confluency and infected with HSV (0.01 MOI) in 10 cm plates. 4 days after infection the supernatant was harvested and centrifuged at 700 RCF for 10 min, then 25-30 mL of supernatant was ultracentrifuged in 5 mL of 20% sucrose in sterile ddH_2_O (overlayed on the supernatant) at 64, 512 RCF for 3h at 4 °C. The supernatant was discarded and the reminder viral solution was resuspended in PBS to a 1:100 ratio of the original supernatant from the Vero cells infection. The virus was then aliquoted and stored at -80 °C. Viral stocks were titled on Vero cells.

### HSV infection of RAW 264.7 cells and BMDM

WT and KO RAW 264.7 cells were seeded (0.45x10^6^ cells/well) in 12 well plates, then infected the next day with 1 mL of DMEM (without serum or P/S) containing HSV KOS (less pathogenetic, yet still infective HSV strain)^63^ (MOI 0.1) and left in the incubator for 1h. After this time, the media was replaced with fresh complete DMEM and cells were returned to the incubator. The cell supernatant was collected 24 and 48h after infection, centrifuged for 15 min at 1, 000 RCF and was used to perform a plaque assay in Vero cells as mentioned above. The resulting WT and KO pellets were used for RT-qPCR to determine the expression of IFNβ mRNA. WT RAW 264.7 (0.25x10^6^ cells/well) were seeded in 12 well plates and treated the day after with 5 μM of Sanofi-14h for 24h. After pre-treatment cells were infected and the plaque assay was performed as described. BMDM were infected in 24 well plates (0.35x10^6^ cells/well) with HSV (MOI 5) in 300 μL of DMEM for 1h. The media then was replaced with regular RPMI without LCCM, cells were returned to the incubator and after 24h the supernatant was harvested, centrifuged at 1, 972 RCF for 15 min and used to performed the plaque assay.

### Sanofi-17a synthesis

Compound Sanofi-17a was synthesized according to the reported procedure using trans-1,4-cyclohexanediol instead of mixtures^31^. The final deprotection of THP was performed using HCl in diethyl ether and dichloromethane to minimize the by-product formation using alcoholic solvents. 1 hNMR spectra of Sanofi-17a matched the reported values.

### RNA sequencing

WT and KO RAW 264.7 cells were plated and treated with DMXAA and Sanofi-14h (0.5 μM) for the indicated times. The RNA was extracted as described and sent to Novogen for sequencing. The details of the strategy used can be found in the supplementary methods.

### RNA Seq analysis

Paired-end RNA-seq data in FASTQ format were mapped to the mouse reference genome, GRCm39 and Gencode mouse vM27 transcript annotation, using the spliced transcripts alignment to a reference (STAR) software^64^. Gene expression estimates as count of mapped sequence reads, and transcripts per million mapped reads (TPM) were estimated using the transcriptome assembler, StringTie2^65^. Differential gene expression analysis was performed using linear models and the Bioconductor R package, limma, and voom normalization, starting from gene expression count^66, 67^. Gene set enrichment analysis were performed on log2 transformed fold-changes (logFC) ranks, with the Bioconductor R package, fgsea^68^. Volcano plots and heatmaps were generated using the R package, ggplot2. ggplot2^69^ and ComplexHeatmap^70^. Batch-corrected TPMs by ComBat method were obtained using the Bioconductor R package, sva^71^, and were used as starting input for the heatmaps. Unless indicated otherwise, the default differentially expressed genes (DEGs) high-lighted in volcano plots and heatmaps were filtered at Benjamini-Hochberg FDR adjusted p-value<0.01, and absolute logFC>1, and average log2 TPM>0.

### Statistical analysis

GraphPad Prism version 9.0.0 for MAC was used to perform One-way ANOVA followed by Dunnett’s multiple comparisons test. Statistical significance was defined by a *P <0.05*. This software was also use to calculate the inhibitors’ IC_50_ values based on the Nonlinear fit function.

## Acknowledgments

We thank the members of the Baldwin and Hagan Lab for their technical and intellectual support. We thank Dr. Matthew McPeek for technical support and Alexia Perryman and Dr. Karel Alcedo for their insightful conversations about this project. This research was supported by the National Institute of Health. Grants: R35-CA197684 and R35-CA197684 to ASB and K08HL143271-01A1 and 1R03HL155249-01 to RSH

---

**Supplemental figures:**
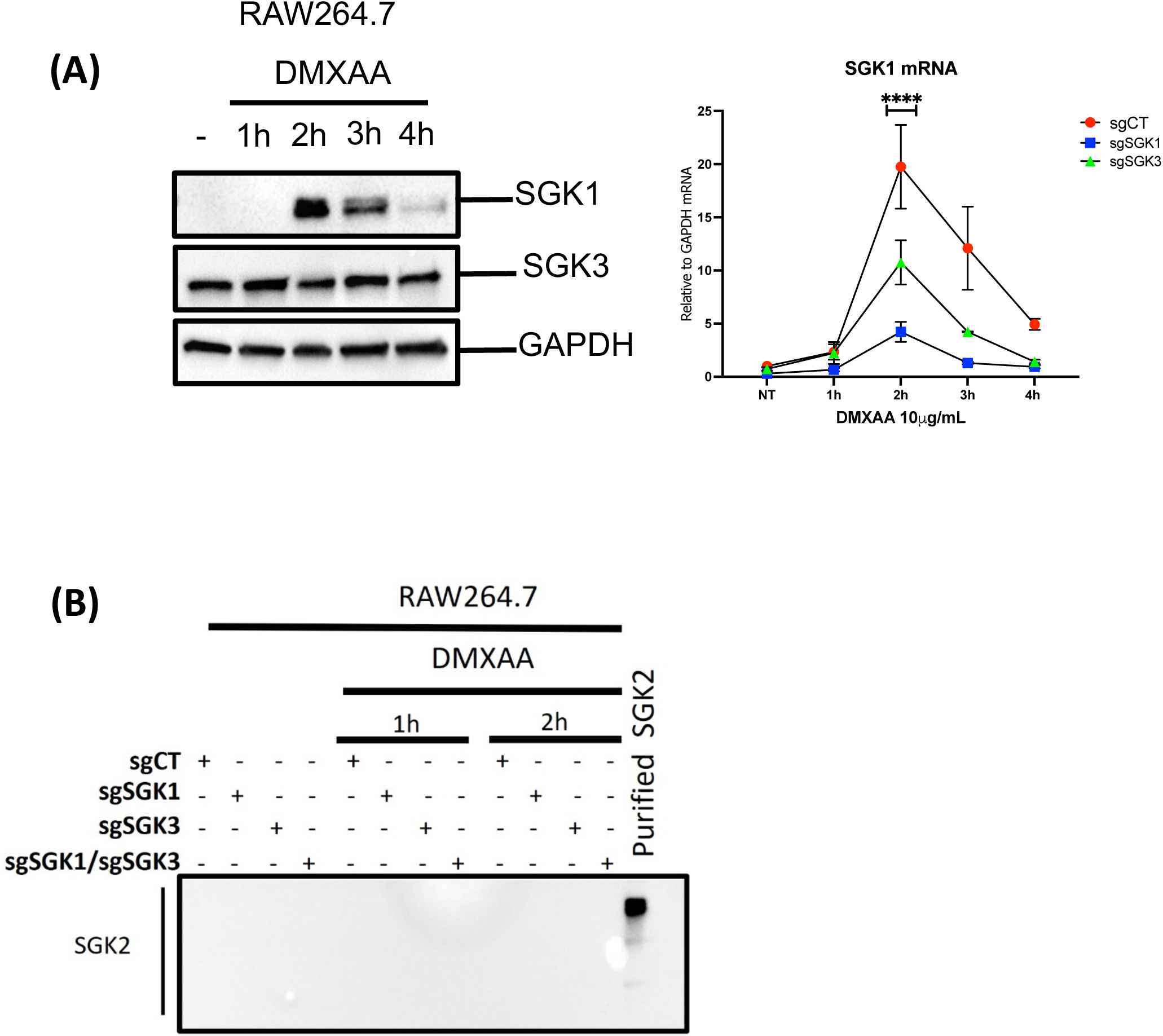
**A)** DMXAA increases mRNA and protein expression of SGK1, but not SGK3 through time. Upper Immunoblot shows the levels of SGK1 protein and left graph depicts the SGK1 mRNA levels after DMXAA treatment at different times in SGK1 and SGK3 single KO cells. **B)** SGK2 is not expressed in RAW264.7 cells at basal or after DMXAA treatment at the protein(immunoblot) or mRNA levels (data not shown).

## Notes

### Competing Interest Statement

The authors have declared no competing interest.

### Summary of Updates

The last name of the corresponding author is Baldwin and not Ernest

